# Bayesian Unidimensional Scaling for visualizing uncertainty in high dimensional datasets with latent ordering of observations

**DOI:** 10.1101/163915

**Authors:** Lan Huong Nguyen, Susan Holmes

## Abstract

**Background:** Detecting patterns in high-dimensional multivariate datasets is non-trivial. Clustering and dimensionality reduction techniques often help in discerning inherent structures. In biological datasets such as microbial community composition or gene expression data, observations can be generated from a continuous process, often unknown. Estimating data points’ ‘natural ordering’ and their corresponding uncertainties can help researchers draw insights about the mechanisms involved.

**Results:** We introduce a Bayesian Unidimensional Scaling (BUDS) technique which extracts dominant sources of variation in high dimensional datasets and produces their visual data summaries, facilitating the exploration of a hidden continuum. The method maps multivariate data points to latent one-dimensional coordinates along their underlying trajectory, and provides estimated uncertainty bounds. By statistically modeling dissimilarities and applying a DiSTATIS registration method to their posterior samples, we are able to incorporate visualizations of uncertainties in the estimated data trajectory across different regions using confidence contours for individual data points. We also illustrate the estimated overall data density across different areas by including density clouds. One-dimensional coordinates recovered by BUDS help researchers discover sample attributes or covariates that are factors driving the main variability in a dataset. We demonstrated usefulness and accuracy of BUDS on a set of published microbiome 16S and RNA-seq and roll call data.

**Conclusions:** Our method effectively recovers and visualizes natural orderings present in datasets. Automatic visualization tools for data exploration and analysis are available at: https://nlhuong.shinyapps.io/visTrajectory/.

## Background

Multivariate, biological data is usually represented in the form of matrices, where features (e.g. genes, species) represent one dimension and observations (e.g. samples, cells) the other. In practice, these matrices have too many features to be visualized without pre-processing. Since a human brain can perceive no more than three dimensions, a large number of methods have been developed to collapse multivariate data to their low-dimensional representations; examples include standard principal component analysis, (PCA), classical, metric and non-metric multidimensional scaling (MDS), as well as more recent diffusion maps, and t-distributed Stochastic Neighbor Embedding (tSNE). While simple 2 and 3D scatter plots of reduced data are visually appealing, alone they do not provide a clear view of what a “natural ordering” of data points should be nor the precision with which such an ordering is known. Continuous processes or gradients often induce “horseshoe” effects in low-dimensional linear projections of datasets involved. Diaconis et al. [1] discuss in detail the horseshoe phenomenon in multidimensional scaling using an example of voting data. Making an assumption that legislators (observations) are uniformly spaced in a latent ideological left-right interval, they showed mathematically why horseshoes are observed. In practice, observations can be collected unevenly along their underlying gradient. Therefore, sampling density differences should be incorporated in an improved model.

In this article, we propose Bayesian Unidimensional Scaling (BUDS), a class of models that maps observations to their latent one-dimensional coordinates and gives measures of uncertainty for the estimated quantities while taking into account varying data density across regions. These coordinates constitute summaries for high-dimensional data vectors, and can be used to explore and to discover the association between the data and various external covariates which might be available. BUDS includes a new statistical model for inter-point dissimilarities, which is used to infer data points’ latent locations. The Bayesian framework allows us to generate visualizations of the estimated uncertainties. Our method produces simple and easy to interpret plots, providing insights to the structure of the data.

### Related work

Recovering data points’ ordering has recently become important in the single cell literature for studying cellular differentiation processes. A number of new algorithms have been proposed to estimate a pseudotemporal ordering, or pseudotime, of cells from their gene expression profiles. Most of the methods are twostage procedures, where the first step is designed to reduce large RNA-seq data to its k-dimensional representation, and the second to recover the ordering of the cells. The second step is performed by computing a minimum spanning tree (MST) or a principal curve on the reduced dataset, [2, 3, 4, 5]. The methods listed are algorithmic and provide only point estimates of the cells’ positions along their transition trajectory.

Very recently, new methods for pseudotime inference have been proposed that incorporate measures of uncertainty. However, they are based on a Gaussian Process Latent Variable Model (GPLVM) [6, 7, 8]. These methods make an assumption, which is not always applicable, that either features or components of a kdimensional projection of the data can be represented by Gaussian Processes. Applying GPLVM to components of reduced data is often more effective as highdimensional biological data tend to be very noisy. For example, Campbell et al. [7] perform pseudotime fitting only on 2D embeddings of gene expression data. Unfortunately, this means that their uncertainty estimates for the inferred pseudotimes do not account for the uncertainties related to the dimensionality reduction step applied previously, and hence might be largely imprecise as the reduced representations might not capture enough of structure in the data. Reid and Wernich [8] on the other hand implemented BGPLVM method directly to the data features (genes), but their method seems practical only when applied to a subset of genes, usually not more than 100. Thus, the method requires the user to choose which features to include in the analysis. Reid and Wernich’s method is semisupervised as it requires capture times, which are a proxy for the latent parameters they want to recover. While this approach is suitable for studying cell states, as it encourages pseudotime estimates to be consistent with capture times, it is not appropriate for fully unsupervised problems, where no prior information on the relative ordering of observations is known.

BUDS models pairwise dissimilarities on the original data directly. This means that BUDS can incorporate information from all features in the data, and can account for all uncertainties involved in the estimation process. Moreover, BUDS is flexible because it gives the user freedom to choose the most suitable dissimilarity metric for the application and type of data under study.

## Methods

In this section, we discuss how we model, analyze and visualize datasets in which a hidden ordering of observations is present. We first give details on our generative Bayesian model and then describe the procedure for constructing visualizations based on the estimated latent variables.

### The model

Biological data is represented as a matrix, *X ∈* ℝ^*p×n*^. Corresponding pairwise dissimilarities, *d*_*ij*_ = *d*(**x**_*i*_, **x**_*j*_), can be computed where **x**_*i*_ *∈* ℝ^*p*^ is an *i*th-column of *X* representing the *i*th-observation. Dissimilarities quantify interactions between observations, and can be used to infer datapoints’ ordering. Since our method targets datasets with latent continua, we can assume that observations within these datasets lie approximately on one-dimensional manifolds (linear or non-linear) embedded in a higher dimensional space. As a result, the inter-point dissimilarities in the original space should be closely related to the distances along the latent data trajectory. Our model recovers the latent positions (1D coordinates) of the datapoints along that unknown trajectory. We take a parametric Bayesian approach and model original dissimilarities as random variables centered at distances in the latent 1D space. This allows us to to draw posterior samples of datapoints’ latent locations. These estimates specify the ordering of the observation according to the most dominant gradient in the data.

The choice of dissimilarity measures in the original space should depend on the type and the distribution of the data. We observed that Jaccard distance seems robust to noise and allows for effective recovery of gradients hidden in 16S rRNA-seq microbial composition data. For gene expression data, we usually use a correlation-based distance, applied to normalized and log-transformed counts, *d*(**x**_*i*_, **x**_*j*_) = (1 − *ρ*(**x**_*i*_, **x**_*j*_))*/*2 where *ρ*(**x**_*i*_, **x**_*j*_) is a Pearson correlation between **x**_*i*_ and **x**_*j*_. On the voting (binary) data we used a kernel *L*_1_ data.

Mathematically, we want to use these pairwise dissimilarities to map high dimensional datapoints, **x**_*i*_, to their latent coordinates, *τ*_*i*_ *∈* [0, 1]. These coordinates represent positions of the observations along their unknown trajectory. The more similar the *i*th and the *j*th observation are, the closer *τ*_*i*_ and *τ*_*j*_ should be. It follows that ***τ*** can also be used for database indexing, where it is of interest to store similar objects closer together for faster lookup. With ***τ*** one can generate many useful visualizations that help understand and discover patterns in the data. To infer ***τ***, we model dissimilarities on high-dimensional data, *d*_*ij*_, as noisy realizations of the true underlying distances *δ*_*ij*_ = *| τ*_*i*_ − *τ*_*j*_*|*.

Overall, our method can be thought of as a special case of Bayesian Multidimensional Scaling technique, whose objective is to recover from a set of pairwise dissimilarities a *k*-dimensional representation of the original data together with associated measures of uncertainty. Previously, Bayesian MDS methods have been implemented in [9, 10], where authors used models based on truncated-normal, and log-normal distributions. These models, however, do not allow for varying levels of noise across different regions of the data. We believe that when modeling dissimilarities one needs to accommodate for heteroscedastic noise, whose scale should be estimated from the data itself. We, thus, developed a model based on a Gamma distribution with a varying scale factor for the noise term:

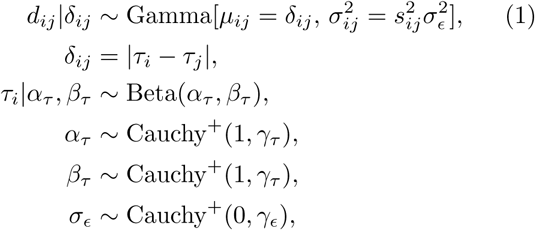

where *s*_*ij*_ *∝ ŝ*(*d*_*ij*_) and *ŝ*^2^(*d*_*ij*_) is an empirical estimate of the variance of *d*_*ij*_ discussed in the next section. Note that the Gamma distribution is usually parametrized with shape and rate (*α, β*) parameters rather than mean and variance (*µ, σ*^2^). The shape and the rate parameter for *d*_*ij*_ can be easily obtained using the following conversion: 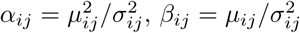. Note that *α*_*τ*_, *β*_*τ*_ are centered around 1, as Beta(1, 1) is similar to the uniform distribution which is the assumed distribution of *τ*_*i*_’s if no prior knowledge of the sampling density is available. However, since *α*_*τ*_ and *β*_*τ*_ are treated as random variables, the BUDS can infer unequal values for the parameters that are away from 1, which means it can model datasets where the sampling density is higher on one or both ends of the data trajectory.

In general, our model postulates a one dimensional gradient along which the true underlying distances are measured with noise. Distances are assumed to have a Gamma distribution, a fairly flexible distribution with a positive support. As dissimilarities are inherently non-negative quantities, Gamma seems to be a reasonable choice. Dissimilarities can be more or less reliable depending on their range and the density or sparsity of the data region, therefore our model incorporates a varying scale factor for the noise term. We estimate the variance of individual *d*_*ij*_’s using the nearest neighbors of the two datapoints **x**_*i*_ and **x**_*j*_ associated with the dissimilarity. The details on how to estimate the scale of the noise term are included in the next section.

Since dissimilarities on high dimensional vectors can have a different range than the ones on 1D coordinates, we incorporate the following shift and scale transformation within our model to bring the distributions of the dissimilarities closer together:

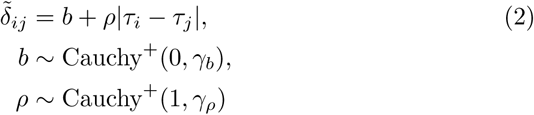

where *b* and *ρ* are treated latent variables, inferred together with ***τ*** from the posterior. Now 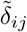 can be substituted for *δ*_*ij*_ in the main model.

In some cases the dissimilarities in high-dimensional settings can be concentrated far away from zero, and provide insufficient contrast between large and small scale interactions between datapoints. The following rank-based transformation can help alleviate the issue and bring the distribution of *d*_*ij*_’s closer to the one of 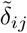’s,

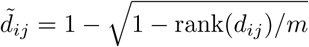

where *m* = *n*(*n* − 1)*/*2 is the number of distinct pairwise dissimilarities, assumed symmetric. The rank-based transformation is similar to techniques used in ordinal Multidimensional Scaling which are aimed at preserving only the ordering of observed dissimilarities [11] and not their values.

### Variance of dissimilarities

Pairwise dissimilarities, either directly observed or computed from the original data can be noisy. The accuracy of dissimilarities in measuring interactions between pairs of observations does not need to be constant across all observations. For example, a dataset might be imbalanced, and some parts of the its latent trajectory might be more densely represented than others. We expect dissimilarities to contain less noise in data-rich regions than in the ones where only a few observations have been collected.

A data-driven approach is taken to estimate scale factors for the noise. We use local information to estimate the variance of individual *d*_*ij*_’s, as illustrated in Fig.. First, for each *d*_*ij*_, we gather a set of *K*-nearest neighbors of **x**_*i*_ and **x**_*j*_, denoted Γ_*K*_(**x**_*i*_) and Γ_*K*_(**x**_*j*_) respectively. We then estimate the variance of *d*_*ij*_ as the empirical variance of distances between **x**_*o*_ and the *K*-nearest-neighbors of **x**_*j*_ and between **x**_*j*_ and the *K*nearest-neighbors of **x**_*i*_. More precisely,

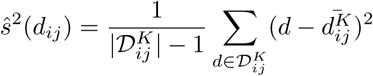

where 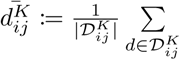 *d* is the average distance over the set 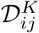 defined as:

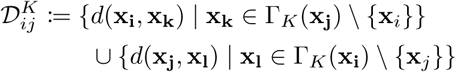

Note that we exclude the **x**_*i*_ from the set of neighbors of **x**_*j*_ and vice versa when gathering the distances in the set 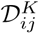. This, is useful in cases when **x**_*i*_ and **x**_*j*_ are within each other’s *K*-nearest-neighborhoods. Without exclusion the set 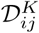 would contain zero-distances *d*(**x**_*i*_, **x**_*i*_) or *d*(**x**_*j*_, **x**_*j*_) which would have an undesirable effect of largely overestimating the variance of *d*_*ij*_.

We use *ŝ*^2^(*d*_*ij*_) only as a relative estimate, and then compute the scale parameter for the noise term as follows:

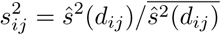

where the bar notation represents the empirical mean over all *ŝ*(*d*_*ij*_)’s. The mean variance for all dissimilarities, 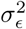 is treated as a latent variable and is estimated together with all other parameters.

The tuning parameter *K* should be set to a small number such that local differences in variances of *d*_*ij*_’s and the data density can be captured. In this paper we used *K* = 10 for all examples, and noticed that the estimates of ***τ*** are robust to different (reasonable) choices of *K*.

### Statistical Inference

Our model is implemented using the STAN probabilistic language for statistical modeling [12]. In particular we use the RStan R package [13] which provides various inference algorithms. In this article we used Automatic Differentiation Variational Inference (ADVI) [14]. ADVI is a “black-box” variational inference program, much faster than automatic inference approaches based on Markov Chain Monte Carlo (MCMC) algorithms. Even though the solutions to variational inference optimization problems are only approximations to the posterior, the algorithm is fast and effective for our applications.

Our model requires a choice of a few hyperparameters *γ*_*τ*_, *γ*_*b*_, *γ*_*ρ*_ and *γ*_*∊*_, which are scale parameters of the half-Cauchy distribution. The half-Cauchy distribution is recommended by Gelman et al. [15, 16] as a weakly informative prior for scale parameters, and a default prior for routine applied use in regression models. It has a broad peak at zero and allows for occasional large coefficients while still performing a reasonable amount of shrinkage for coefficients near the central value [16]. The scale hyperparameters were set at 2.5, as we do not expect very large deviations from the mean values. The value 2.5 is also recommended by Gelman in [16], and is a default choice for positive scale parameters in many models described in the RStan software manual [13].

### Visual representations of data ordering

We developed visual tools for inferring and studying patterns related to the natural ordering in the data. Our visualizations uncover hidden trajectories with corresponding uncertainties. They also show how sampling density varies along a latent curve, i.e. how well a dataset covers different regions of an underlying gradient. We implemented a multi-view design with a set of visual components: 1) a plot of latent ***τ*** against its ranking, 2) a plot of ***τ*** against a sample covariate, 3) a heatmap of reordered data, 4) a 2D and a 3D posterior trajectory plot, 5) a data density plot, 7) a datapoint location confidence contour plot, 8) a feature curves plot. The settings panel and the visualization interface are depicted in Fig. 2 and 3).

**Figure 1.**
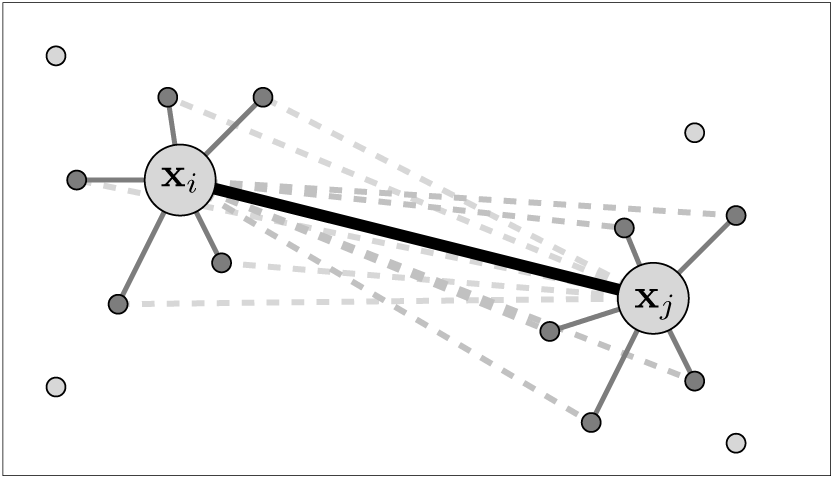
Graphical representation of points **x**_*i*_ and **x**_*j*_ together with their neighbors. The set of (dashed) distances from **x**_*i*_ to the *K*-nearest-neighbors of **x**_*j*_, and from **x**_*i*_ to the *K*-nearest-neighbors of **x**_*i*_ is used to compute *ŝ*^2^(*d*_*ij*_), the estimate of the variance of *d*_*ij*_. Here we chose *K* = 5.

**Figure 2.**
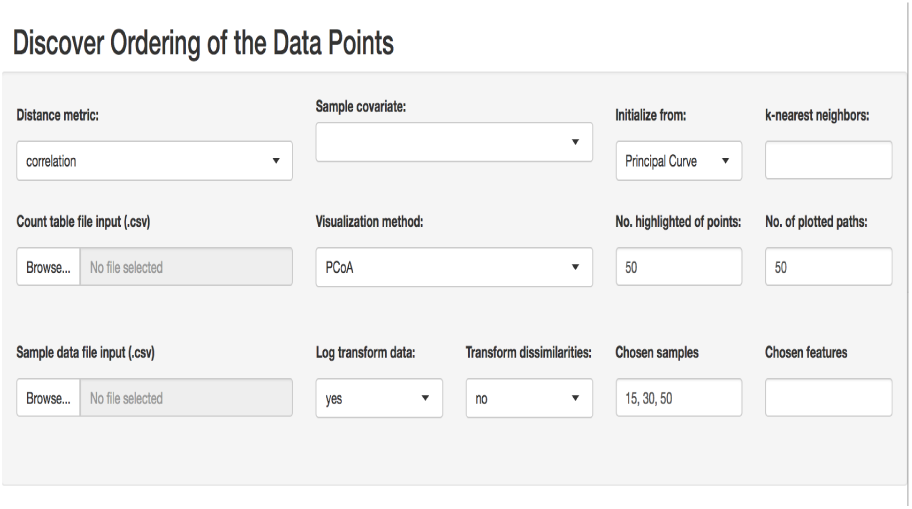
The settings panel for BUDS visualization interface, where the data and the supplementary sample covariates can be uploaded, and specific features and samples as well as other parameters can be selected.

**Figure 3.**
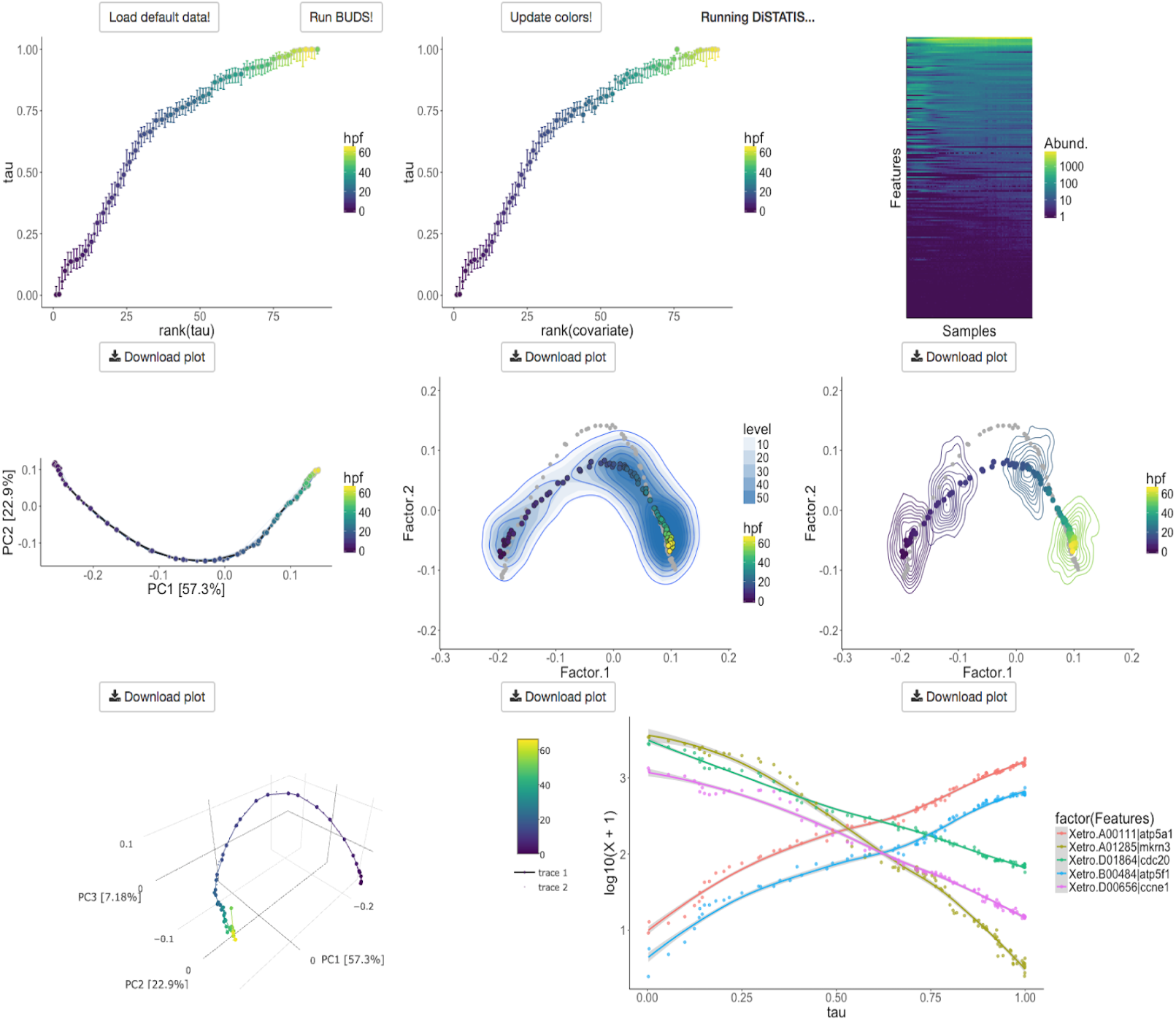
The visualization interface for latent data ordering. The plots are shown for the frog embryo gene expression dataset collected by Owens et al. [20]. First row, left to right: a plot of ***τ*** against its ranking, a plot of ***τ*** against a sample covariate, a heatmap of reordered data. Second row, left to right: 2D posterior trajectory plot, data density plot, datapoint location condition confidence contour plot. Third row: 3D data trajectory plot and features curves plot.

For our visualizations we chose a recently developed `viridis` color map, designed analytically to “perfectly perceptually-uniform” as well as color-blind friendly [17]. This color palette is effective for heatmaps and other visualizations and has now been implemented as a default choice in many visualization packages such as `plotly` [18] or `heatmaply` [19].

#### Latent ordering plot view

The most direct way to explore the ordering in the data is through a scatter plot of ***τ***-coordinates against their ranking (Fig. 4), which includes measures of uncertainty. This view depicts the arrangement of the datapoints along their hidden trajectory, and shows how confident we are in their estimated location along the path.

**Figure 4.**
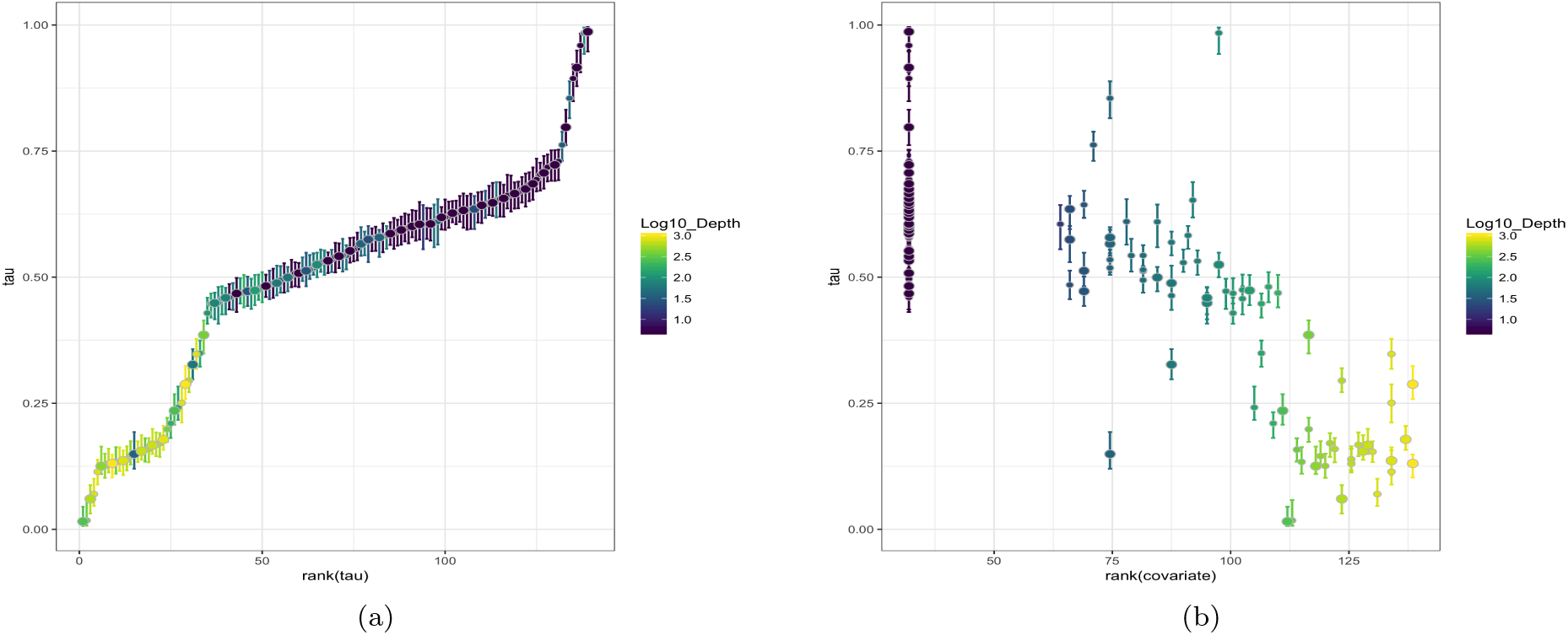
Latent ordering in TAℝA Oceans dataset shown with uncertainties. The differences in the slope of plot (a) indicate varying data coverage along the underlying gradient. Correlation between the water depth and the latent ordering in microbial composition data is shown in (b). Coloring corresponds to log_10_ of the water depth (in meters) at which the ocean sample was collected.

Despite its simple form, the plot provides useful information about the data. For example, the variability in the slope of the plot indicates how well covered the corresponding region of the trajectory is, i.e. if the slope is steep and the value of ***τ*** changes faster than its rank, then the data is sparse in that region.

Color-coding can be also used to explore which sample attributes are associated with the natural ordering recovered, e.g. in the results section we show that samples ordering is associated with the water depth in TARA Oceans dataset, and with the age of the infant in the DIABIMMUNE dataset. One can also plot the estimates of ***τ*** against the ranking of a selected covariate to examine the correlation between the two more directly. If a high correlation is observed, one might further test whether the feature is indeed an important factor driving the main source of variability in the data. For example, when analyzing microbiome data one might discover environmental or clinical factors differentiating collected samples.

#### Reordered heatmaps

Heatmaps are frequently used to visualize data matrices, *X*. It is a common practice to perform hierarchical clustering on features and observations (rows and columns of *X*) to group similar objects together and allow hidden block patterns to emerge. However, clustering only suggests which rows and columns to bundle together; it does not provide information on how to order them within a group. Thus, it is not a straightforward matter how to arrange features and samples in a heatmap. Matrix reordering techniques, sometimes called seriation, are discussed in a comprehensive review by Innar Liiv [21]. Many software programs are also available for generating heatmaps, among which NeatMap [22] is a non-clustering one, designed to respect data topology, and intended for genomic data. Here we describe our own take on matrix reordering for heatmaps.

As our method deals with situations which involve a continuous process rather than discrete classes, hierarchical clustering approach for organizing data matrices might be suboptimal. Instead, since our method estimates ***τ***, which corresponds to a natural ordering of the samples, we can use it to arrange the columns of a heatmap. To reorder the rows of *X*, we use an inner product between features and ***τ***. More specifically, we compute a vector, **z ∈** ℝ^*p*^, whose elements are dot products equal to 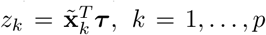 where 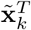 is the *k*th row of 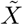, a (column) normalized data matrix. The dot product is used for row-ordering as it reflects the mean *τ* value for the corresponding feature. In other words *z*_*k*_ indicates the mean location along the latent trajectory (expected *τ*) where feature *k* “resides”. Using this value for ordering seems natural, as the features with similar “*τ*-utility” levels are placed close together. Fig. 5 shows example heatmaps produced by our procedure when applied to real microbial data.

**Figure 5.**
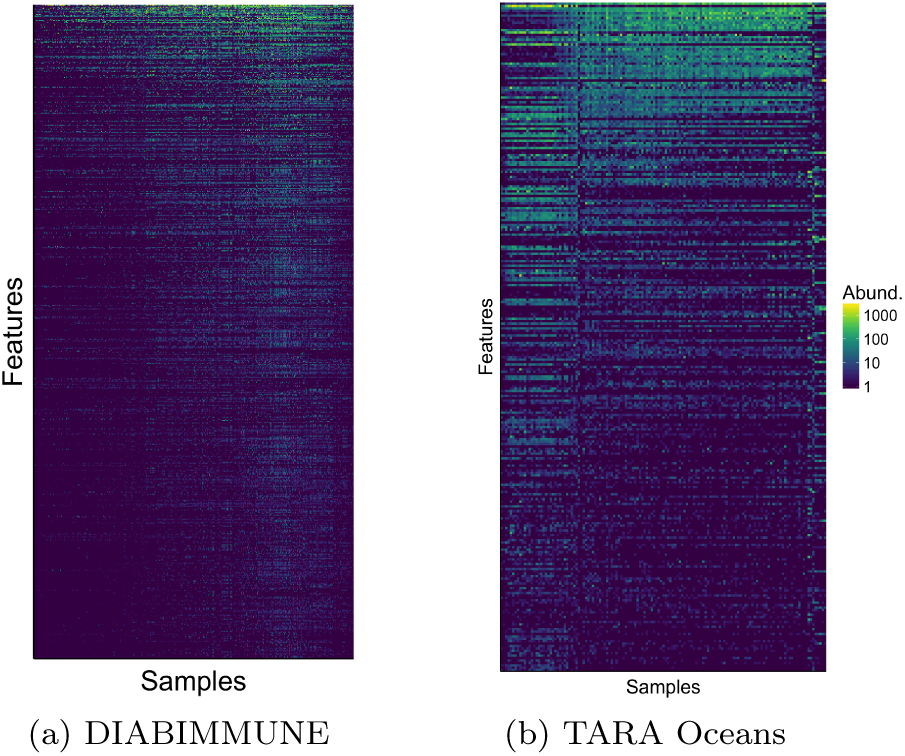
Reordered heatmaps for two microbial composition datasets.

We include the comparison of our heatmaps with the ones generated by NeatMaps in Additional file 1, Fig. S1, S2. NeatMaps computes an ordering for rows and columns of an input data matrix, however it does not provide any uncertainty estimates. Moreover, the method is not optimized for cases where the data lies on a non-linear 1D manifold. BUDS heatmaps often display smooth data structures such as banded or triangular matrices. These, structures help users discover which groups of variables (genes, species or other features) have higher or lower values in which part of the underlying dominant gradient in the data.

#### Feature dynamics

The ordering of observations can also be used to explore the trends in variability of individual data features with respect to the discovered latent gradient. For example, when studying cell development processes, we might be interested in changes of expression of particular genes. The expression levels plotted against the pseudo-temporal ordering estimated with ***τ*** along with the corresponding smoothing curves, can provide insights into when specific genes are switched on and off (see Fig. 6 (a)). Similarly, when analyzing microbiome data one can find out which species are more or less abundant at which regions of the underlying continuous gradient (see Fig. 6 (b)). Both of the plots are discussed more in detail in the results section.

**Figure 6.**
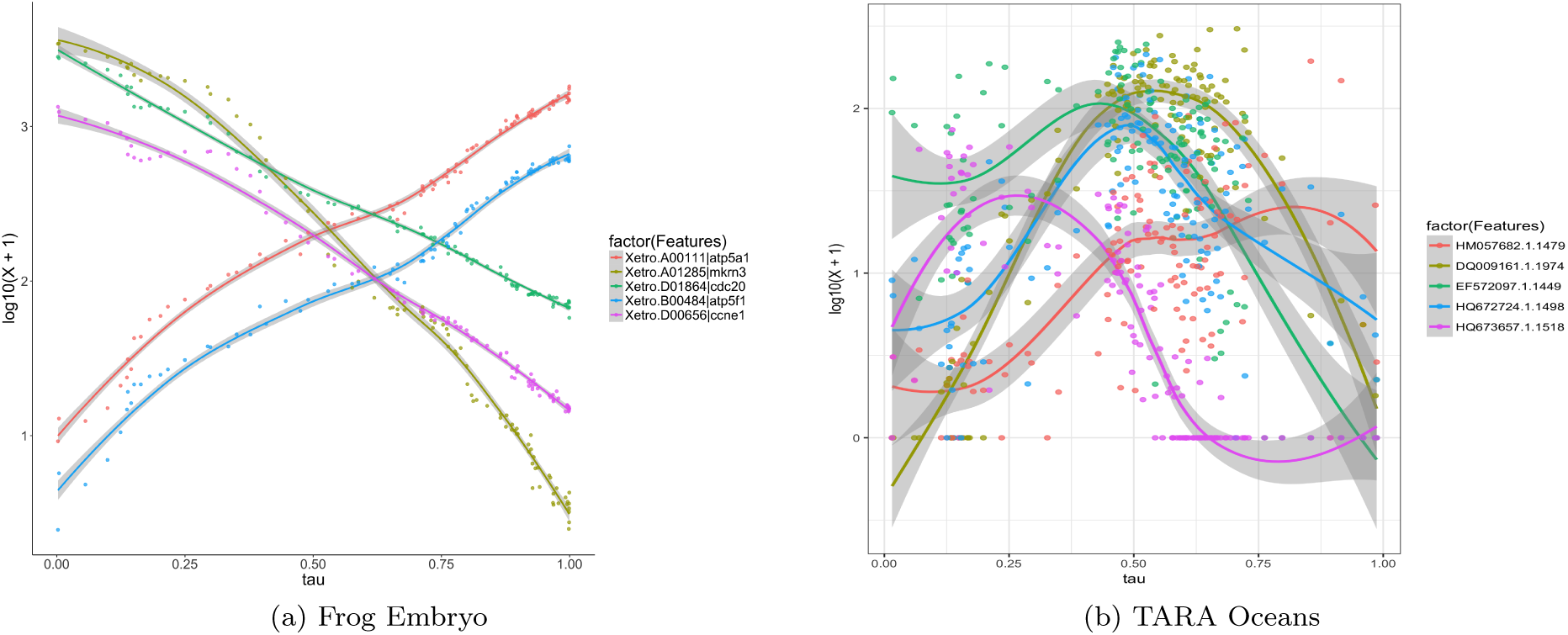
Feature dynamics along inferred data trajectory. Frog embryo gene expression levels follow smooth trends over time (a). Selected bacteria are more abundant in TAℝA Oceans samples corresponding to the right end of the latent interval representing deeper ocean waters (b).

#### Trajectory plots

Often it is also useful to visualize a data trajectory in 2 or 3D. We use dimensional reduction methods such as principal coordinate analysis (PCoA) and t-distributed stochastic neighbor embedding (tSNE) [23] on computed dissimilarities to display lowdimensional representations of the data. PCoA is a linear projection method, while t-SNE is a non-convex and non-linear technique.

After plotting datapoints in the reduced two or three dimensional space, we superimpose the estimated trajectories, i.e. we add paths which connect observations according to the ordering specified by posterior samples of ***τ***. We usually show 50 posterior trajectories (see blue lines in Fig. 7), and one highlighted trajectory that corresponds to the posterior mode-***τ*** estimate. To avoid a crowded view, the mode-trajectory is shown as a path connecting only a subset of points evenly distributed along the gradient, i.e. corresponding to *τ*_*i*_’s evenly spaced in [0, 1]. We also include a 3D plot of the trajectory as the first two principal axes sometimes do not explain all the variability of interest. The third axis might provide additional information. The 3D plot provides an interactive view of the ‘mode-trajectory’; it allows the user to rotate the graph to obtain an optimal view of the estimated path. The rotation feature also facilitates generating 2D plots with any combination of two out of three principal components (0PC1, PC2, PC3), which is an efficient alternative to including three separate plots.

**Figure 7.**
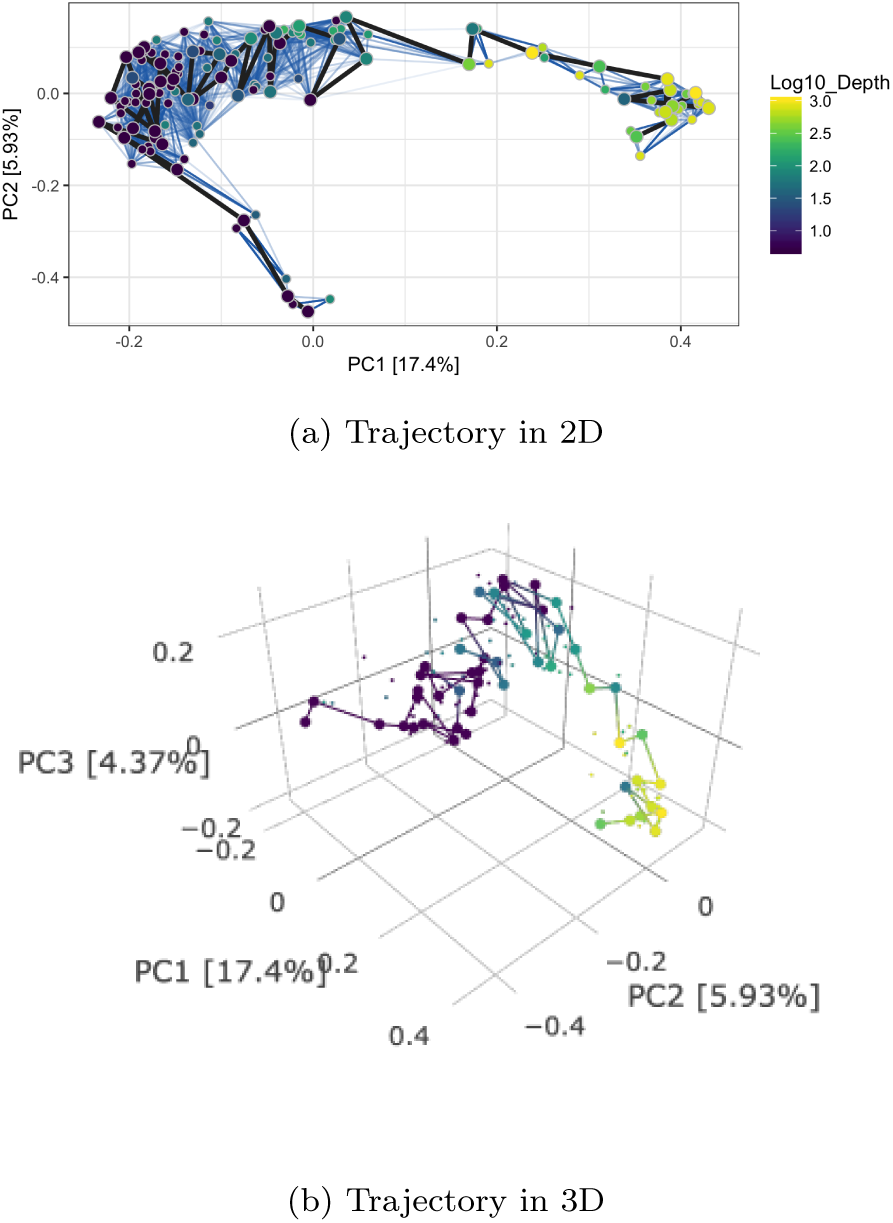
Posterior trajectory plots for TARA Oceans dataset; 50 paths are plotted in blue to show uncertainties in the inferred ordering. “Mode”-trajectory is shown in black for a subset of highlighted (bigger) datapoints evenly spaced along the *τ −*interval (a). The same mode-trajectory is show in a 3D (b). The axis are labeled by the principal component index and the corresponding percent variance explained.

#### Data density and uncertainty plots

Since it might be also of interest to visualize data density along its trajectory, we also provide 2D plots with density clouds. We use posterior samples of ***τ*** ^*\**^ to obtain copies of latent distance matrices, 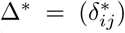. Then we generate copies of noisy dissimilarity matrices, *D*^*\**^, by drawing from a Gamma distribution centered at elements of Δ^*\**^ according to our model. We combine posterior dissimilarity matrices in a data cube (*T ∈* ℝ^*n×n×t*^) where *t* is the number of posterior samples, and then apply DiSTATIS, a three-way metric multidimensional scaling method, [24] to obtain data projections together with their uncertainty and density estimates.

Our visualization interface includes two plots displaying the data configuration computed with DiSTATIS. The first depicts the overall data density across the regions, and the second shows confidence contours for selected individual datapoints. Contour lines and color shading are commonly used for visualizing non-parametrically estimated multivariate-data density [25, 26]. Contour lines, also called isolines, joint points of equal data density estimated from the frequency of the datapoints across the studied region. Apart from visualizing data density, we use isolines also to display our confidence in the estimated datapoints’ locations. BUDS can generate posterior draws of dissimilarity matrices which are used by DiSTATIS to obtain posterior samples of data coordinates in 2D. These coordinates are used for non-parametric estimation of the probability density function (pdf) of the true underlying position of an individual observation. Contour lines are then used delineate levels of the estimated pdf. These contours have a similar interpretation as the 1D error bars included in *τ*-scatter-plots and visualize the reliability of our estimates

Fig. 8 (a) shows density clouds for DiSTATIS projections and the consensus points representing the center of agreement between coordinates estimated from each posterior dissimilarity matrix *D*^*\**^. From the density plot one can read which regions of the trajectory are denser or sparser than the others. Fig. 8 (b) gives an example of a contour plot for four selected datapoints using TARA Oceans dataset discussed in the results section. The size of the contours indicate the confidence we have in the location estimate, the large the area covered by the isolines, the less confident we are in the position of the observation.

**Figure 8.**
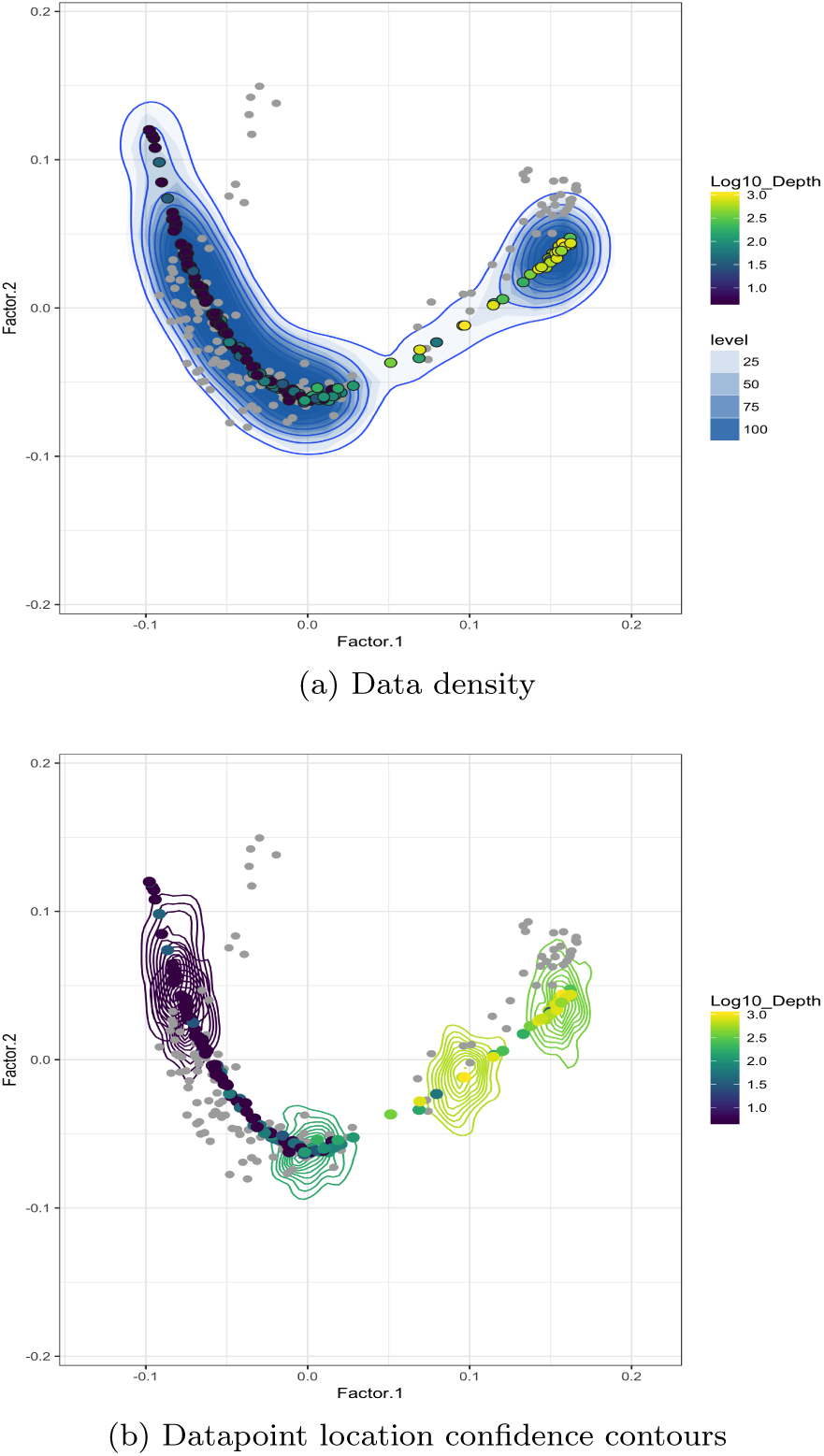
Overall data density plot (a) and confidence contours for location estimates of selected datapoints (b) for TARA Oceans dataset. Colored points denote DiSTATIS consensus coordinates, and gray ones the original data.

Our visualization interface is implemented as a Shiny Application [27] and 3D plots were rendered using the Plotly R library[18].

## Results

We demonstrate the effectiveness of our modeling and visualization methods on four publicly available datasets. Bayesian Unidimensional Scaling is applied to uncover trajectories of samples in two microbial 16S, one gene expression and one roll call dataset.

### Frog embryo gene expression

In this section we demonstrate BUDS performance on gene expression data from a study on transcript kinetics during embryonic development by Owens et al. [20]. Transcript kinetics is the rate of change in transcript abundance with time. The dataset has been collected at a very high temporal resolution, and clearly displays the dynamics of gene expression levels, which makes it well suited for testing the effectiveness of our method in detecting and recovering continuous gradient effects.

The authors of this study collected eggs of Xenopus tropicalis and obtained their gene expression profiles at 30-min intervals for the first 24 hrs after an in vitro fertilization, and then hourly sampling up to 66 hr (90 samples in total). The data was collected for two clutches; here we only analyze samples from Clutch A for which poly(A)+ RNA was sequenced. For our analysis, we used the published transcript per million (TPM) data, from which we remove the ERCC spike-ins and rescale accordingly. A log base 10 transformation (plus a pseudocount of one) is then applied to avoid heteroscedasticity related to high variability in noise levels among genes with different mean expression levels. This is a common practice for variance stabilization of RNA-seq data. As inputs to BUDS, we used Pearson correlation based dissimilarities defined in the methods section.

As shown in Fig. 3 and 9, BUDS accurately recovered the temporal ordering of the samples using only the dissimilarities computed on the log-expression data. We also observe that samples collected in the later half have more similar gene expression profiles, than the ones sequenced immediately post fertilization, as BUDS tend to place the samples sequenced 30+ hours after fertilization (HPF) closer together on the latent interval, than the ones from the first 24h. In other words gene expressions undergo rapid changes in the early stages of the frog embryonic development, and slow down later on.

**Figure 9.**
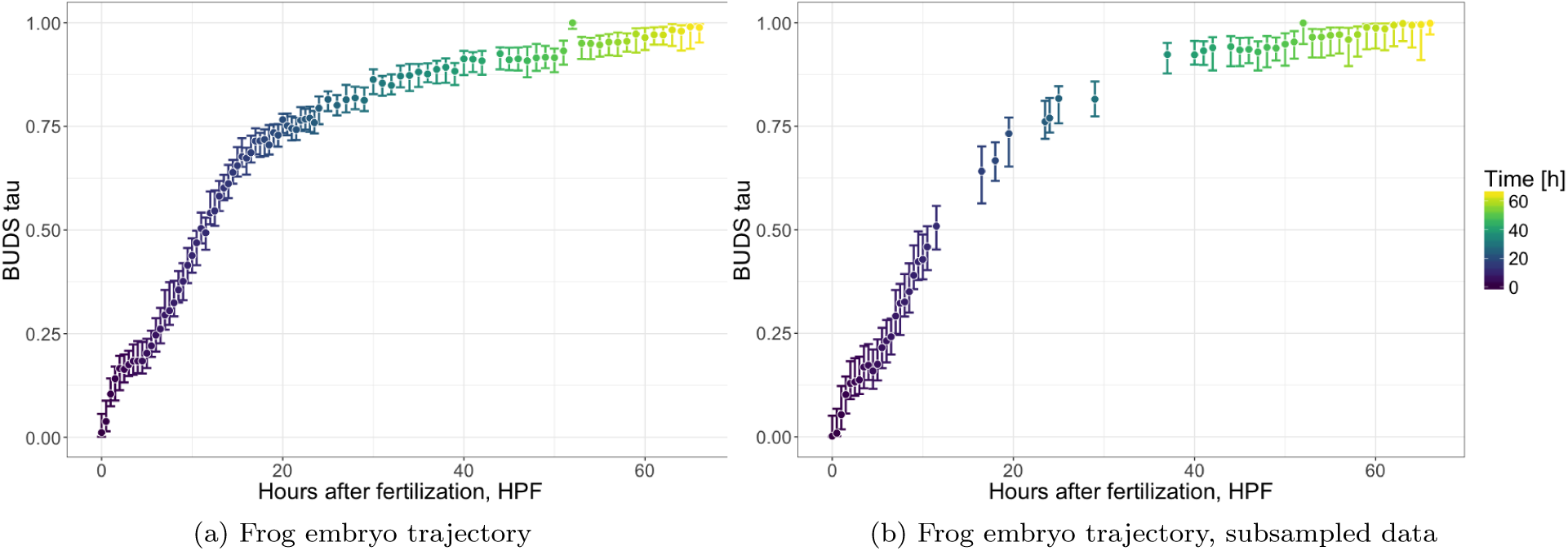
Frog embryonic development trajectory. Correlation between inferred location on the latent trajectory and time in hours post fertilization (HPF). Latent coordinates computed using BUDS on untransformed Pearson correlation distances.

To show that our method is robust to differences in sampling density along data trajectory, we subsampled the dataset keeping only every fourth datapoint from the period between 10 and 40 HPF, and all samples from the remaining time periods. We observed that the samples’ ordering recovered stays consistent with the actual time (in terms of HPF). As desired, the 95%-HPDI are wider in sparser regions, i.e. samples 24hr+ after fertilization, as they were collected in 1hr intervals instead of 30-min. In particular, the downsampled region [10-40 HPF], involves estimates with clearly larger uncertainty bounds.

Additionally, the visual interface was used to show the trends in expression levels of selected individual genes. Fig. 6 depicts how expression levels of five selected genes vary along the recovered data trajectory, which in this case corresponds to time post fertilization. We can see that the expression drops for three genes and increases for two others.

### TARA Oceans microbiome

The second microbial dataset comes from a study by Sunagawa et al. [28] conducted as a part of TARA Oceans expedition, whose aim was to characterize the structure of the global ocean microbiome; 139 samples were collected from 68 locations in epipelagic and mesopelagic waters across the globe. Illumina DNA shotgun metagenomic sequencing was performed on each prokaryote-enriched samples, and the taxonomic profiles were obtained from merged reads (miTAGs) that contained signatures of 16S rRNA gene, which were then mapped to the sequences in the SILVA reference database and clustered into 97% OTUs. The paper reported that the ocean microbiome composition and diversity was stratified according to the depth at which the samples were collected rather than the geographical location. Microbial composition was thus found to be ordered along a gradient associated with the water depth.

BUDS was used to estimate a natural ordering of samples from Jaccard pairwise distances computed on bacterial composition data. Fig. 4 shows resulting *τ* coordinates plots. We detected a high correlation of the latent coordinates with the depth of water at which a sample was collected (Spearman rank correlation *ρ ≈* 0.79). This result is consistent with the trends observed in the original study. In Fig. 4 (a), we can see that the samples were not collected evenly along their trajectory. Some intervals such as *τ ∈* [0.75, 1.0] and *τ ∈* [0.2, 0.45] have a much steeper slope than others indicating sparser sampling in the corresponding region. A similar conclusion can be reached by looking at the 2D density plot, Fig. 7 (a)), where certain parts of TARA Oceans data trajectory are much sparser than the rest. Identifying sparsities along the data trajectory might be of interest to researchers, who might like to determine whether the sparsity stems from the absolute rarity of certain type of samples or from the flaws of the data collection procedure which include physical restrictions (e.g. the ocean stations were located at specific depth). We observed that all 63 surface water layer samples had been recorded water depth of 5m, which suggests that either the depth measurements were not be very precise, or the study by design did not cover a variety (in terms of water depth) of surface samples. In Fig. 7 (b) the surface samples are all grouped in a single column on the left side; however the BUDS estimates of their location still suggests some biological variability within the group. Collecting more observations in sparse regions researchers can gain a more comprehensive understanding of the underlying process and potentially discover the cause of the rarity.

### Infant gut microbiome

Under the DIABIMMUNE project, Kostic et al. [29] studied the dynamics of the human gut microbiome in its early stage and the progression toward type 1 diabetes (T1D). Their cohort consisted of 33 infants from Finland and Estonia, genetically predisposed to T1D. Children were followed from birth until 3 years of age, and their stool samples were collected monthly by their parents. Amplicon sequencing was done on the V4 region of the 16S rDNA and taxonomic profiling performed with QIIME [30]. Additionally, Kostic et al. performed functional analysis using shotgun metagenomic sequencing. Here we are only interested in the progression of the gut microbiome to ‘maturity’, and want to estimate its developmental trajectory. Similarly to the approach taken in section of the original paper discussing microbial community dynamics, we exclude T1D cases. To further reduce the biological variability related to factors other than time, we limit our analysis to the 14 infants in the control group from Finland, who are in the same T1D HLA risk group (HLA = 3, the biggest risk group), leaving 327 samples in total.

We analyzed the bacterial composition dynamics using the published operational taxonomic unit (OTU) count table. We filtered OTU at prevalence level of 5 (present in ≥ 5 samples) and total count of 300 (sum in all samples ≥ 300), leaving 1341 OTUs. As we wanted to show change in the bacterial composition related to new bacterial species acquisition, we used as input Jaccard distances which use species presence/absence information.

We showed that the gut microbiome follows a clear developmental gradient, and that the ordering of samples according to the posterior mode of ***τ*** is correlated with the infant’s age at time of collection (with Spearman rank correlation *ρ ≈* 0.74). Moreover, we observed that the location estimates were more variable for samples collected early after birth, see Fig. 10. For example, notice that the first 100 samples (corresponding to the earliest period after birth) take up the same space on the latent ***τ*** [0, 1]-interval as more than 200 remaining ones. This indicates that observations collected later in the study are more similar to each other than the ones gathered right after birth.

**Figure 10.**
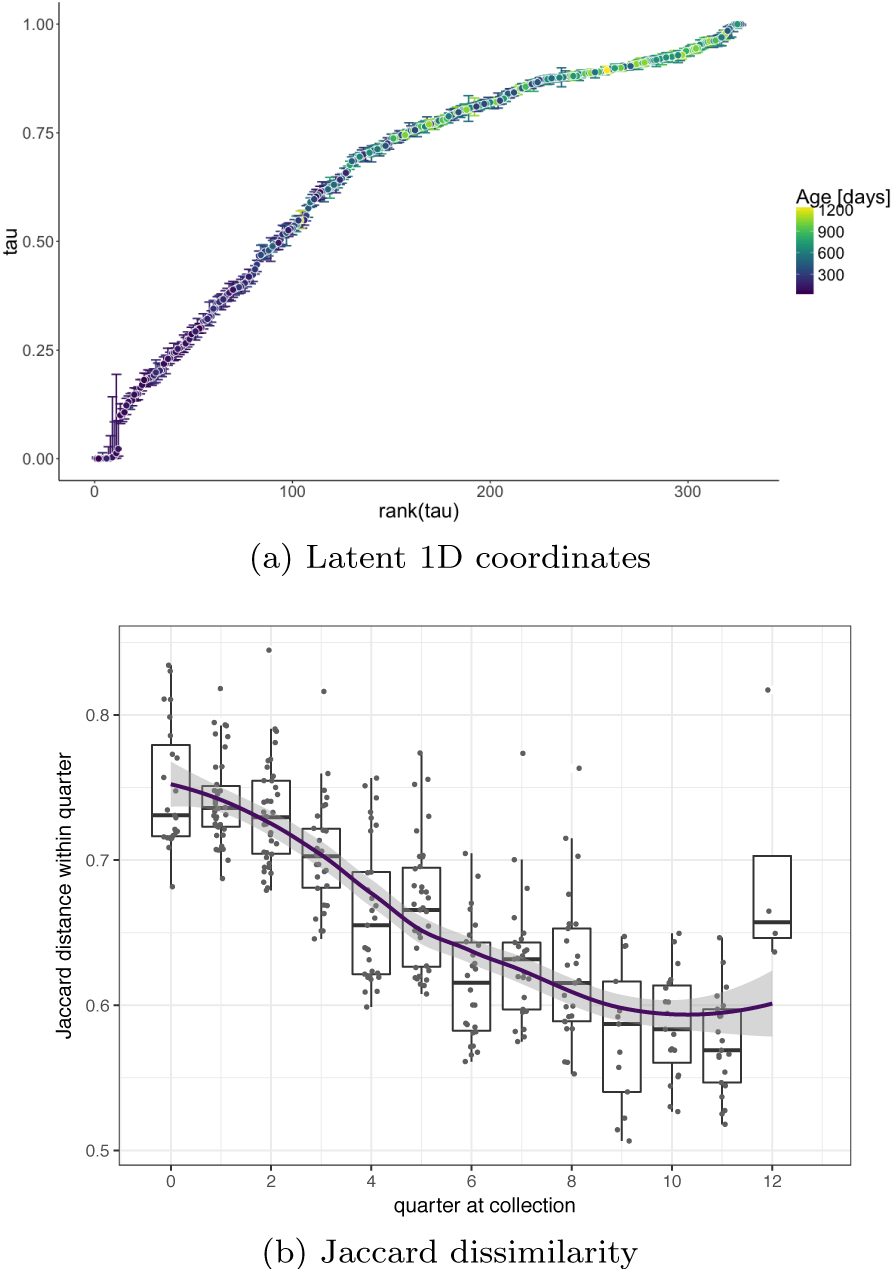
Gut microbiome development trajectory. Latent 1D coordinates along the estimated BUDS trajectory shown with colored intervals indicating the 95% highest posterior density intervals (HPDI) (a), Jaccard dissimilarity between samples within each quarter after birth (b).

To confirm our hypothesis that the bacterial composition is more similar among samples collected later in the study than right after birth, we computed and compared the average Jaccard distance between each sample pairs collected in the same quarter (across all individuals). As shown in Fig. 10(b), Jaccard dissimilarity displays a decreasing trend over time. Samples from the first few quarters just after birth are more variable than the ones collected closer to the end of the study. This agrees with our guess that bacterial composition seems to progress to a ‘mature’ gut microbiome state, and that while after birth the gut microflora might vary a lot across subjects, infants seem to acquire a shared ‘microbial base’ during the first three years.

Additionally, looking at the reordered data heatmap, we see a triangular structure appearing. This suggests that while there are some bacteria, that are present in the gut microbiome from the beginning or appear early after birth, there are other which are acquire later in the infant’s life. This pattern is visible in Fig. 5 (a) where the top rows show bacteria which are present almost in every samples, while the bottom rows represent taxa which appear in the infant gut in later stages corresponding to the right side of the heatmap.

### Roll Call Data

So far we have presented only biological data examples. However, our method is general enough not to be limited to a specific type of data. To show this, we applied BUDS to voting data similar to Diaconis et al. in [1]. We obtained the 114th U.S. Congress Roll Call data containing votes of 100 senators on a total of 502 bills within the period between January 3, 2015 and January 3, 2017. We computed a kernel *L*_1_ distance on the binary “yea” vs “nay” data, and applied BUDS to estimate the ordering of the senators related to their voting pattens.

Fig. 11 shows the ordering obtained with our procedure. We see that without any knowledge of senators’ party membership and only using the binary voting data, BUDS was able to separate Democrats from Republicans on a left-right political spectrum. It is interesting that senator Bernie Sanders was ranked in this ordering as the most liberal one (the first on the left). This seems to be plausible, given his proposed policies during 2016 U.S. election campaign. Overall, the ordering computed by BUDS seems to reflect senators’ optimal “ideological utility” position, which is a rather abstract concept. The accuracy of our results could be evaluated further using outside information and domain knowledge.

**Figure 11.**
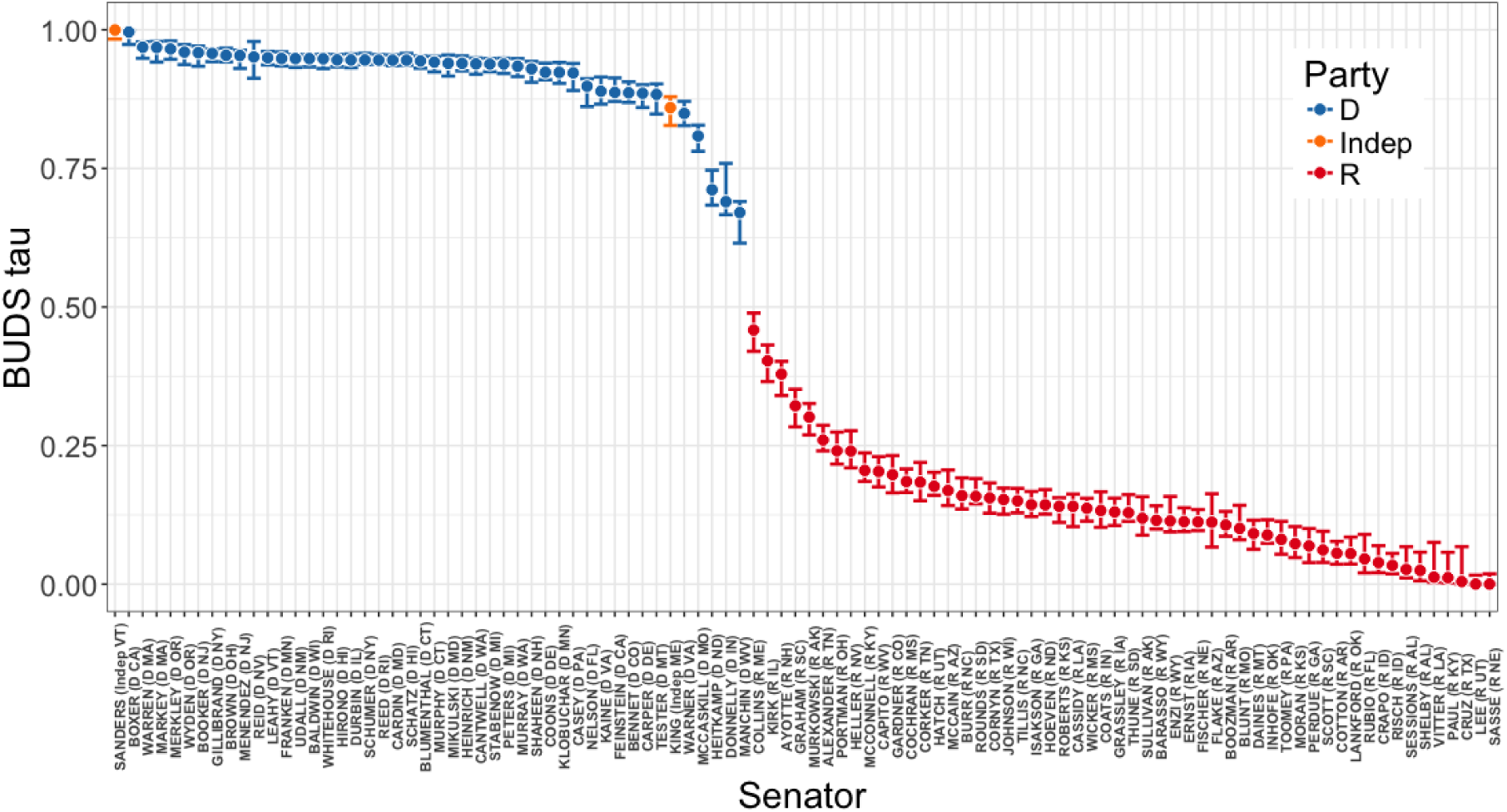
Ordering of 100 Senators based on 114th U.S. Congress ℝoll Call data.

## Discussion and Conclusions

In biology, we often encounter situations where a gradual change or a progression between data points can be observed. The analysis of datasets originating from these processes requires different sets of tools than standard machine learning, and pattern recognition techniques such as simple clustering. Here we presented our new visualization interface developed specifically for exploring datasets which involve a latent continuum. We designed visual representations of the data which clearly illustrate the natural ordering of data points. While tools exist which support the tasks, existing approaches are not comprehensive, lack measures of uncertainty or involve unrealistic assumptions.

Our novel modeling approach is based on a Bayesian framework, model observed inter-point dissimilarities with latent distances on a one-dimensional interval representing the underlying data trajectory. Thus, it is highly different from the current pseudotime inference methods which rely on an assumption that either the features or the low-dimensional representation components of the data come from a Gaussian process. We impose no such restrictions, and only require that the observed dissimilarities are noisy realizations of distances between latent coordinates in a one-dimensional space. This is a reasonable condition, as it is our objective to find an ordering of observations that respects the interactions observed in the original data.

Using the estimated natural arrangement of observations, our program generates a set of plots that facilitate understanding of an underlying process that shapes the data. We include plots of the latent coordinates, reordered heatmaps, smooth curve plots showing dynamics of individual features (e.g. specific gene expression or species abundance) as well as 2 and 3D visualizations of the data which incorporate measures of uncertainty.

We demonstrated the effectiveness of our visualizations on publicly available datasets. Our plots showed that the infant gut microbiome changes with time throughout the first three years after birth, which is consistent with the findings of the original study by Kostic et al. [29]. Moreover, by inspecting the plot of the inferred latent coordinates against its ranking we found that the bacterial composition stabilizes, after rapid changes during the first year of life. The estimated coordinates represent a pseudo-temporal ordering, and provide a better ordering than the real time, as the gut bacterial flora develops at various rates within different infants. Similar trends were observed on the frog embryo development dataset from a study by Owens et al. [20]. For TARA Oceans dataset published by Sunagawa et al. [28] we showed that the collected samples do not cover the microbial composition gradient evenly. Some types of ocean samples were more sparsely represented than others. Our method was also successful at mapping senators on the leftright political spectrum based on 114th U.S. Congress voting data. BUDS has clearly separated Democrats from Republicans without any knowledge of the legislators’ party affiliations.

In conclusion, we have developed a useful tool for analyzing datasets in which latent orderings occur. Our program is automatic and user friendly. Using our interactive visualization interface, researchers can generate illustrative plots which facilitate the process of data exploration and understanding.

## List of abbreviations used

BUDS: Bayesian Unidimensional Scaling
MDS: Multidimensional Scaling
tSNE: t-distributed Stochastic Neighbor Embedding
HPDI: Highest Posterior Density Interval

## Declarations

**Availability of data and materials**

**BUDS Rstan package** for fitting and drawing posteriors from the specified Bayesian model: https://github.com/nlhuong/buds.

**visTrajectory Shiny App** for generating associated visualizations:

https://github.com/nlhuong/visTrajectory.

Additionally, a web-based application, not requiring any installations, is available at https://nlhuong.shinyapps.io/visTrajectory/.

**Instructional demo video** explaining BUDS’ interactive visualization interface: https://github.com/nlhuong/visTrajectory/tree/master/video_demo.

## Authors’ contributions

LHN designed, developed and implemented the method, as well as applied it to the data examples provided. SH gave the general idea of the problem, and provided theoretical and practical guidance. All the authors read and approved the final manuscript.

## Ethics approval and consent to participate

Not applicable.

## Consent for publication

Not applicable.

## Author details

^1^ Institute for Computational and Mathematical Engineering, Stanford University, 94305 Stanford, US. ^2^Department of Statistics, Stanford University, 94305 Stanford, US.

## Additional Files

Additional file 1 — Supplementary figures

---

**Figure S1.**
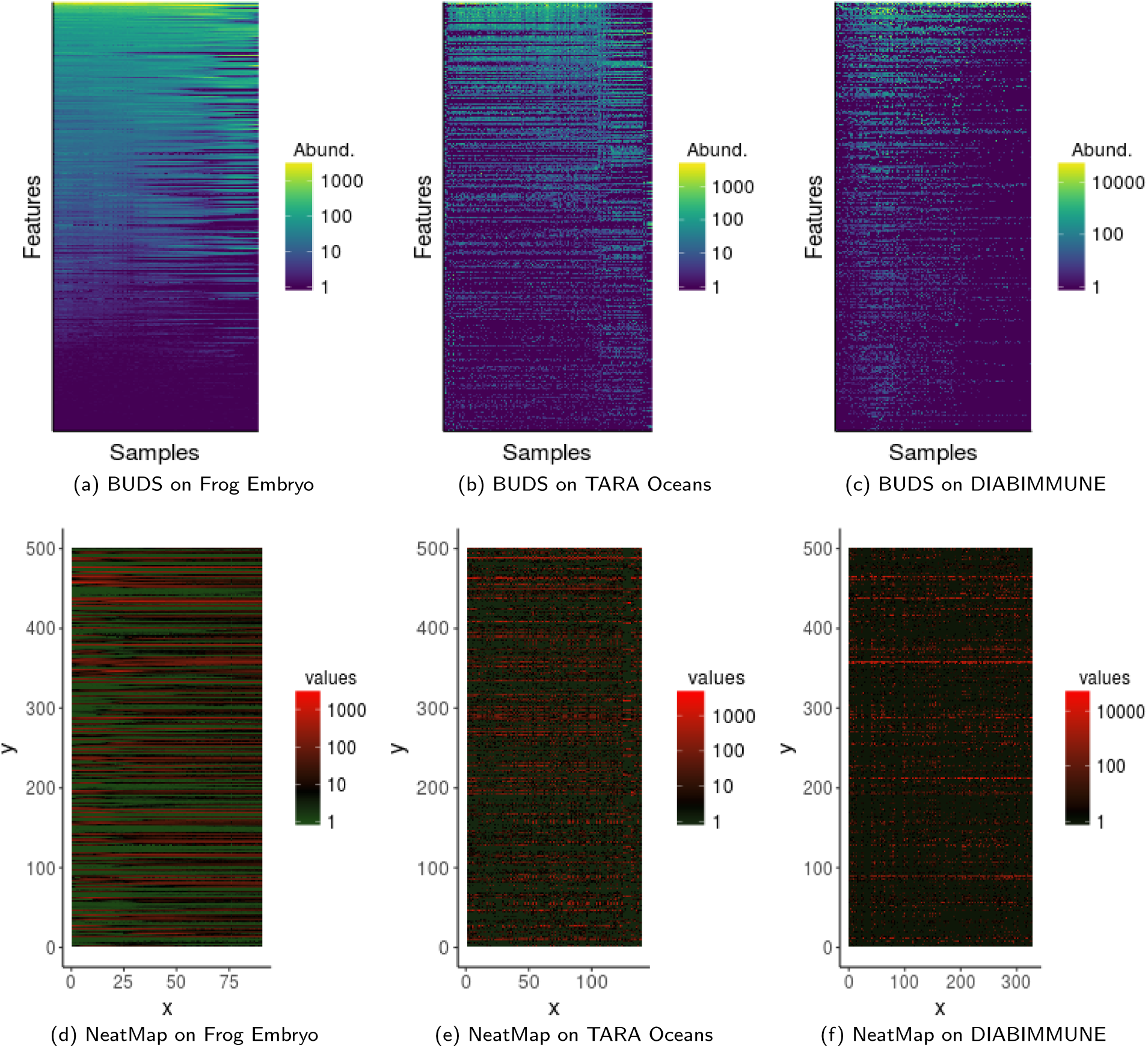
Comparison of BUDS and NeatMap for matrix ordering applied to three biological datasets. The heatmaps are shown for 500 randomly selected features (the same for BUDS and Neatmap). A default color scheme setting was for NeatMap heatmaps. BUDS ordering gives a much clearer visualization of the continuous patterns present in the data.

**Figure S2.**
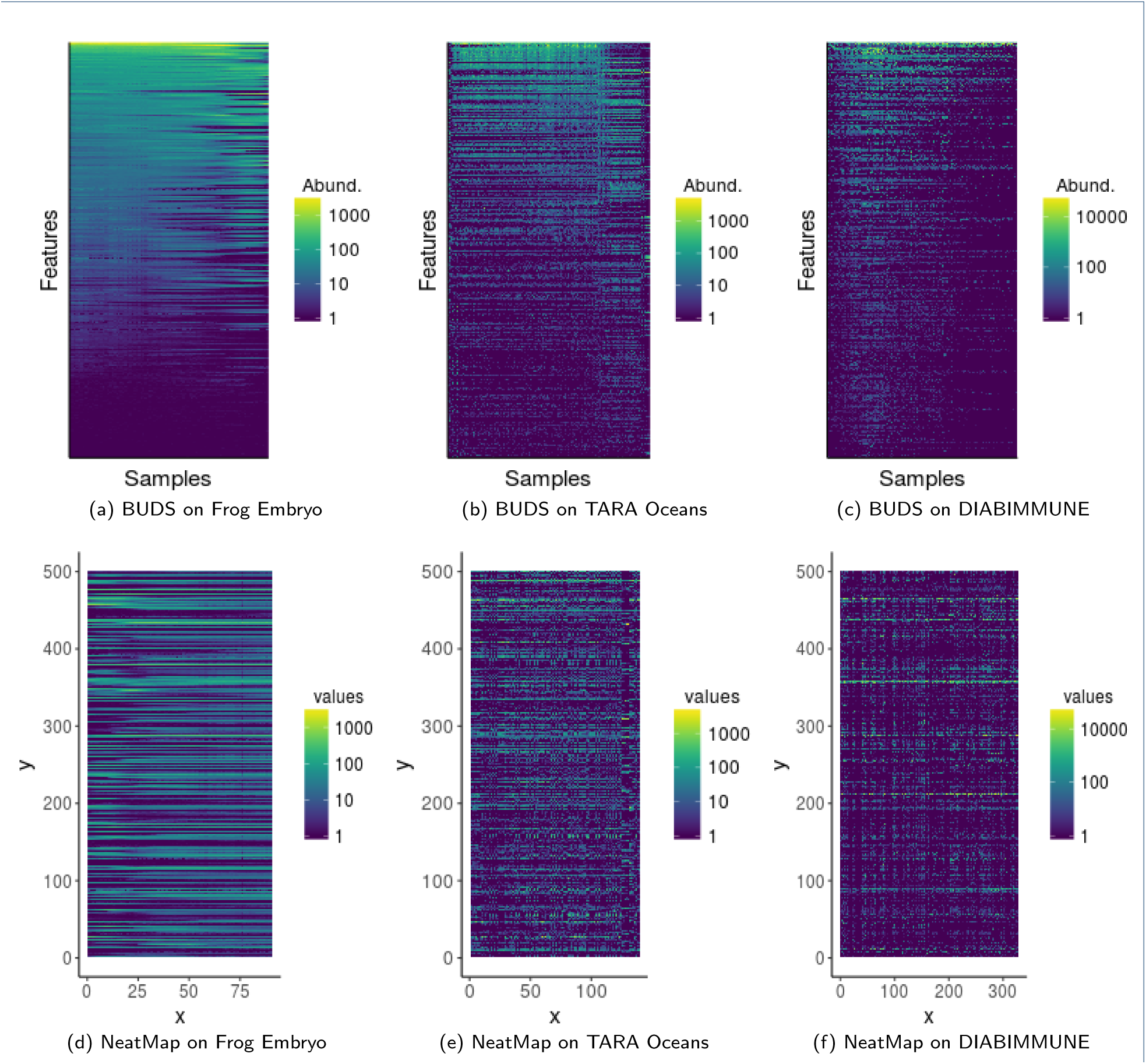
Comparison of BUDS and NeatMap for matrix ordering applied to three biological datasets. The heatmaps are shown for 500 randomly selected features (the same for BUDS and Neatmap). A viridis color scheme setting was for NeatMap heatmaps. BUDS ordering gives a much clearer visualization of the continuous patterns present in the data.

## References

1. Diaconis, P., Goel, S., Holmes, S.: Horseshoes in multidimensional scaling and local kernel methods. Ann. Appl. Stat. 2(3), 777–807 (2008)

2. Trapnell, C., Cacchiarelli, D., Grimsby, J., Pokharel, P., Li, S., Morse, M., Lennon, N.J., Livak, K.J., Mikkelsen, T.S., Rinn, J.L.: The dynamics and regulators of cell fate decisions are revealed by pseudotemporal ordering of single cells. Nat Biotech 32(4), 381–386 (2014). Research

3. Ji, Z., Ji, H.: TSCAN: Pseudo-time reconstruction and evaluation in single-cell RNA-seq analysis. Nucleic Acids Research (2016)

4. Shin, J., Berg, D.A., Zhu, Y., Shin, J.Y., Song, J., Bonaguidi, M.A., Enikolopov, G., Nauen, D.W., Christian, K.M., Ming, G.-l., Song, H.: Single-Cell RNA-Seq with Waterfall Reveals Molecular Cascades underlying Adult Neurogenesis. Cell Stem Cell 17(3), 360–372 (2015)

5. Petropoulos, S., Edsgard, D., Reinius, B., Deng, Q., Panula, S., Codeluppi, S., Reyes, A.P., Linnarsson, S., Sandberg, R., Lanner, F.: Single-Cell RNA-Seq Reveals Lineage and X Chromosome Dynamics in Human Preimplantation Embryos. Cell 165(4), 1012–1026 (2016)

6. Campbell, K., Yau, C.: Bayesian Gaussian Process Latent Variable Models for pseudotime inference in single-cell RNA-seq data. bioRxiv (2015)

7. Campbell, K.R., Yau, C.: Order Under Uncertainty: Robust Differential Expression Analysis Using Probabilistic Models for Pseudotime Inference. PLOS Computational Biology 12(11), 1–20 (2016)

8. Reid, J.E., Wernisch, L.: Pseudotime estimation: deconfounding single cell time series. Bioinformatics 32(19), 2973 (2016)

9. Oh, M.-S., Raftery, A.E.: Bayesian Multidimensional Scaling and Choice of Dimension. Journal of the American Statistical Association 96(455), 1031–1044 (2001)

10. Bakker, R., Poole, K.T.: Bayesian Metric Multidimensional Scaling. Political Analysis 21(1), 125 (2013)

11. Borg, I., Groenen, P.J.F.: Modern Multidimensional Scaling: Theory and Applications, 1st edn. Springer series in statistics. Springer, United States (1997)

12. Carpenter, B., Gelman, A., Hoffman, M., Lee, D., Goodrich, B., Betancourt, M., Brubaker, M., Guo, J., Li, P., Riddell, A.: Stan: A Probabilistic Programming Language. Journal of Statistical Software 76(1), 1–32 (2017)

13. Stan Development Team: RStan: the R interface to Stan. R package version 2.14.1 (2016). http://mc-stan.org/

14. Kucukelbir, A., Ranganath, R., Gelman, A., Blei, D.M.: Automatic Variational Inference in Stan. In: Proceedings of the 28th International Conference on Neural Information Processing Systems. NIPS’15, pp. 568–576. MIT Press, Cambridge, MA, USA (2015)

15. Gelman, A.: Prior distributions for variance parameters in hierarchical models (comment on article by Browne and Draper). Bayesian Anal. 1(3), 515–534 (2006)

16. Gelman, A., Jakulin, A., Pittau, M.G., Su, Y.-S.: A weakly informative default prior distribution for logistic and other regression models. Ann.Appl. Stat. 2(4), 1360–1383 (2008)

17. Garnier, S.: viridis: Default Color Maps from ‘matplotlib’. (2016). R package version 0.3.4. https://CRAN.R-project.org/package=viridis

18. Sievert, C., Parmer, C., Hocking, T., Chamberlain, S., Ram, K., Corvellec, M., Despouy, P.: plotly: Create Interactive Web Graphics Via ‘plotly.js’. (2016). R package version 4.5.6.https://CRAN.R-project.org/package=plotly

19. Galili, T.: heatmaply: Interactive Cluster Heat Maps Using ‘plotly’. (2017). R package version 0.10.1.https://CRAN.R-project.org/package=heatmaply

20. Owens, N.D.L., Blitz, I.L., Lane, M.A., Patrushev, I., Overton, J.D., Gilchrist, M.J., Cho, K.W.Y., Khokha, M.K.: Measuring Absolute RNA Copy Numbers at High Temporal Resolution Reveals Transcriptome Kinetics in Development. Cell Reports 14(3), 632–647 (2016)

21. Liiv, I.: Seriation and matrix reordering methods: An historical overview. Statistical Analysis and Data Mining 3(2), 70–91 (2010)

22. Rajaram, S., Oono, Y.: NeatMap - non-clustering heat map alternatives in R. BMC Bioinformatics 11(1), 45 (2010)

23. van der Maaten, L.J.P., Hinton, G.E.: Visualizing high-dimensional data using t-sne. Journal of Machine Learning Research 9, 2579–2605 (2008)

24. Abdi, H., Williams, L.J., Valentin, D., Bennani-Dosse, M.: STATIS and DISTATIS: optimum multitable principal component analysis and three way metric multidimensional scaling. Wiley Interdisciplinary Reviews: Computational Statistics 4(2), 124–167 (2012)

25. Scott, D.W., Sain, S.R.: Multidimensional Density Estimation. Handbook of Statistics 24, 229–261 (2005)

26. Scott, D.W.: In: Gentle, J.E., Härdle, W.K., Mori, Y. (eds.) Multivariate Density Estimation and Visualization, pp. 549–569. Springer, Berlin, Heidelberg (2012)

27. Chang, W., Cheng, J., Allaire, J., Xie, Y., McPherson, J.: shiny: Web Application Framework for R. (2017). R package version 1.0.3. https://CRAN.R-project.org/package=shiny

28. Sunagawa, S., Coelho, L.P., Chaffron, S., Kultima, J.R., Labadie, K.,Salazar, G., Djahanschiri, B., Zeller, G., Mende, D.R., Alberti, A., Cornejo-Castillo, F.M., Costea, P.I., Cruaud, C., d’Ovidio, F., Engelen, S., Ferrera, I., Gasol, J.M., Guidi, L., Hildebrand, F., Kokoszka, F., Lepoivre, C., Lima-Mendez, G., Poulain, J., Poulos, B.T.,Royo-Llonch, M., Sarmento, H., Vieira-Silva, S., Dimier, C., Picheral, M., Searson, S., Kandels-Lewis, S.,, Bowler, C., de Vargas, C., Gorsky, G., Grimsley, N., Hingamp, P., Iudicone, D., Jaillon, O., Not, F., Ogata, H., Pesant, S., Speich, S., Stemmann, L., Sullivan, M.B., Weissenbach, J., Wincker, P., Karsenti, E., Raes, J., Acinas, S.G., Bork, P.: Structure and function of the global ocean microbiome. Science 348(6237) (2015)

29. Kostic, A., Gevers, D., Siljander, H., Vatanen, T., Hyotylainen, T., Hamalainen, A.-M., Peet, A., Tillmann, V., Poho, P., Mattila, I., Lahdesmaki, H., Franzosa, E.A., Vaarala, O., de Goffau, M., Harmsen, H., Ilonen, J., Virtanen, S.M., Clish, C.B., Oresic, M., Huttenhower, C., Knip, M., Xavier, R.J.: The Dynamics of the Human Infant Gut Microbiome in Development and in Progression toward Type 1 Diabetes. Cell Host & Microbe 17(2), 260–273 (2016)

30. Caporaso, J.G., Kuczynski, J., Stombaugh, J., Bittinger, K., Bushman, F.D., Costello, E.K., Fierer, N., Pena, A.G., Goodrich, J.K., Gordon, J.I., Huttley, G.A., Kelley, S.T., Knights, D., Koenig, J.E., Ley, R.E., Lozupone, C.A., McDonald, D., Muegge, B.D., Pirrung, M., Reeder, J., Sevinsky, J.R., Turnbaugh, P.J., Walters, W.A., Widmann, J., Yatsunenko, T., Zaneveld, J., Knight, R.: QIIME allows analysis of high-throughput community sequencing data. Nat Meth 7(5), 335–336 (2010)

